# From Generation to Discrimination: Vision Foundation Models for Synthetic SEM Image Detection

**DOI:** 10.64898/2026.08.12.744545

**Authors:** A. Palangattu, A. K. Sah, S. Raman, K. Pushpavanam

**Affiliations:** Department of Chemical Engineering, IIT Gandhinagar, Gujarat, India, 382355; ShieldWorkz OT Systems Pvt. Ltd., Benniganahalli, Bengaluru, Karnataka, India, 560093; Department of Computer Science and Engineering, IIT Gandhinagar, Gujarat, India, 382355

**Author notes:** To whom all correspondence must be addressed Shanmuganathan Raman, Ph.D., Department of Computer Science and Engineering Indian Institute of Technology Gandhinagar Gujarat, 382355, India, Karthik Pushpavanam, Ph.D. Department of Chemical Engineering, Indian Institute of Technology Gandhinagar Gujarat, 382355, India.

**Keywords:** Scanning Electron Microscopy, Generative Adversarial Networks, Adaptive Discriminator Augmentation, Contrastive Language-Image Pre-training, Vision Transformer

## Abstract

In materials science, the integrity of scanning electron microscopy (SEM) images is paramount for quality control and validation of research outcomes. However, the introduction of sophisticated generative artificial intelligence, particularly Generative Adversarial Networks (GANs), has introduced a novel vulnerability: the potential for highly realistic, artificially synthesized SEM images to be used fraudulently in scientific literature. To address this challenge, we present a deep learning-based framework capable of distinguishing between authentic SEM images and those synthesized by Generative Adversarial Networks (GANs). Using FastGAN and StyleGAN2-ADA, two state-of-the-art GAN models, we generated synthetic SEM datasets to complement real imaging data. We fine-tuned a pre-trained Contrastive Language-Image Pre-training (CLIP) Vision Transformer (ViT-L-14) for binary classification. By unfreezing the final transformer blocks and appending a custom classification head, the model effectively captures the subtle, high-level artifacts inherent in GAN-generated upsampling. This work highlights the potential of deep learning to safeguard scientific imaging workflows and provides an important step toward detecting and mitigating image forgeries in materials science publications.

## INTRODUCTION

Counterfeiting presents a significant obstacle across various sectors, from financial systems and luxury markets to the digital landscape and identity verification processes.^1,2^ It has been a persistent challenge throughout history, impacting economies and industries worldwide. The circulation of counterfeit currency, for instance, has long troubled governments and financial institutions due to its potential to erode trust in monetary systems and facilitate illicit activities.^3^ Similarly, the production of counterfeit luxury goods not only violates intellectual property rights but also deceives consumers and undermines legitimate businesses, posing significant ethical and economic concerns.^2^

In the digital era, counterfeiting has evolved significantly, driven by the widespread use of digital media and the accessibility of technology to the general population.^1^ The emergence of deepfakes, which leverage AI and machine learning algorithms to generate convincing yet fabricated audio and video content, is of particular concern.^4^ These sophisticated manipulations raise questions about media authenticity and trustworthiness, with far-reaching implications across sectors such as politics, entertainment, and cybersecurity.^3,4^ Recent advancements in image synthesis techniques, such as diffusion models and GANs, have sparked public concern, raising fears that discerning real from fake imagery is becoming increasingly challenging.^5^ Therefore, the ability to discriminate between the real and fake becomes crucial.

Scanning Electron Microscopy (SEM) is a vital tool in the physical and materials sciences, enabling meticulous examination of surface topography, material integrity, and nanoscale structures. Industries ranging from semiconductor manufacturing to aerospace heavily rely on the precision of these images to maintain strict quality standards.^6,7^ Consequently, the authenticity of SEM imagery is a foundational pillar of trust in both industrial quality control and academic publications.^8^

The integrity of scientific imaging has been increasingly threatened recently by cases of picture forgery and modification in research publications.^8,9^ A large-scale analysis of published biomedical papers found that approximately 3.8% contained inappropriate image duplication, with at least half showing features suggestive of deliberate manipulation.^10^ The rapid development of image synthesis methods such as Generative Adversarial Networks (GANs) poses a risk of worsening this problem. With the ability to learn a dataset’s underlying distribution, modern unconditional GANs can create highly convincing images that are often undetectable through manual inspection or metadata analysis. The appearance of fake SEM images in published work risks misleading downstream research, wasting resources, and undermining safety.^8,9^

To distinguish between authentic SEM images and synthetic images generated by FastGAN and StyleGAN2-ADA, we employed a vision foundation model based on a Vision Transformer (ViT) architecture.^11^ Traditional Convolutional Neural Networks (CNNs) extract hierarchical image features through convolutional filters with localized receptive fields and have achieved remarkable success in numerous computer vision tasks.^12,13^ However, their emphasis on local feature extraction may limit their ability to directly model long-range spatial relationships within an image.^11,12,13^ In contrast, Vision Transformer (ViT)-based models employ self-attention mechanisms that capture global contextual information more effectively.^11^ Specifically, we adapt the Contrastive Language-Image Pre-training (CLIP) model, using a Vision Transformer (ViT-L-14) backbone.^11,14^ By fine-tuning the terminal transformer blocks of the CLIP architecture, our model accurately captures nuanced discrepancies between authentic SEM images and those generated by state-of-the-art architectures such as FastGAN and StyleGAN2-ADA, across nanoparticle and biological SEM datasets. FastGAN incorporates a Skip-Layer Channel-wise Excitation (SLE) module, while StyleGAN2-ADA uses adaptive discriminator augmentation, with both architectures designed to improve generative performance on relatively small datasets, making them suitable for SEM datasets where large quantities of annotated images are generally unavailable.^15,16^

The generated images were quantitatively assessed using Mean Squared Error (MSE),^17^ Learned Perceptual Image Patch Similarity (LPIPS),^14^ and Fréchet Inception Distance (FID)^18^ between the synthetic images and their corresponding real SEM images. To ensure transparency in the model’s predictive behaviour, we utilize EigenCAM visualizations, which provide qualitative insight into the spatial regions associated with strong activations in the classifier’s learned feature representations.^19^ Through systematic evaluation across two SEM datasets and two GAN architectures, we assess the ability of a CLIP-based vision foundation model to distinguish AI-generated SEM images from authentic ones and examine the factors underlying its predictions.

## METHODOLOGY

### Dataset and Preprocessing

The proposed framework was evaluated using two independent scanning electron microscope (SEM) image datasets representing distinct application domains: nanoparticle and biological SEM images. Employing two morphologically different datasets enabled assessment of the proposed detector both within individual SEM domains and across domains through cross-dataset evaluation.

Prior to GAN training, all images underwent identical preprocessing. Metadata and scale bars present at the image boundaries were removed through 1:1 cropping to eliminate non-structural information that could bias the classifier. The cropped images were subsequently resized to 256×256 pixels, providing a uniform image resolution for both GAN training and classifier development while preserving the relevant microstructural features.

To construct the “fake” class, synthetic SEM images were generated independently for each dataset using two state-of-the-art unconditional GANs: FastGAN and StyleGAN2-ADA. Separate GAN models were trained for the nanoparticle and biological datasets to preserve the distinct structural characteristics of each imaging domain. Half of the synthetic images in each dataset were generated with FastGAN, and the remaining half with StyleGAN2-ADA, resulting in a balanced collection of GAN-generated SEM images.

### Synthetic Dataset Generation

The official PyTorch implementations of FastGAN and StyleGAN2-ADA were obtained from their respective public repositories and trained from scratch using independently compiled nanoparticle and biological SEM datasets.^15,16^ All images used for training are of 256×256 pixels, and both models were trained from scratch. Unless otherwise specified, the default settings of the respective implementations were retained.

Training was executed independently for each model on a single NVIDIA V100 GPU according to the following experimental parameters:

● **FastGAN Framework:** The generator and discriminator networks were initialized with 64 baseline channels, paired with a latent vector dimension of 256. Dynamic data augmentation was introduced via random horizontal flips.^15^ The architecture was optimized using a batch size of 16 over a total regimen of 100,000 iterations.
● **StyleGAN2-ADA Framework:** To maximize morphological diversity from limited inputs, the data execution pipeline incorporated image mirroring in addition to the native adaptive augmentation loop.^16^ The model was optimized with a batch size of 16 over 1,000 training ticks.

### Quantitative Analysis

Three complementary quantitative metrics, namely Mean Squared Error (MSE), Learned Perceptual Image Patch Similarity (LPIPS), and Fréchet Inception Distance (FID), were employed to evaluate the quality of the generated SEM images.

Mean Squared Error (MSE) was used to quantify the pixel-wise differences between real and generated SEM images. Each real image was compared with all generated images from the corresponding dataset as defined below.

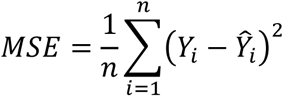

Here *n* is the number of data points, *Y_i_* the observed value, and *Ŷ_i_* the predicted value.^17^

Learned Perceptual Image Patch Similarity (LPIPS) was used to quantify perceptual differences between real and generated SEM images^14^. The AlexNet-based LPIPS implementation was used for the analysis.^14,20^ For each real image, the LPIPS distance was calculated against all generated images, similar to the MSE calculations. This procedure identified the generated image with the smallest perceptual distance from each real image, and the resulting distributions were used for quantitative comparison.

Fréchet Inception Distance (FID) was used to quantify the similarity between the distributions of real and generated SEM images.^18^ FID was calculated using the Torch-Fidelity implementation with the default Inception-based feature extractor.^18,21^ The real and generated image sets were independently processed to obtain their feature representations, from which the mean feature vectors and covariance matrices were calculated. The Fréchet distance between the two resulting feature distributions was then computed as,

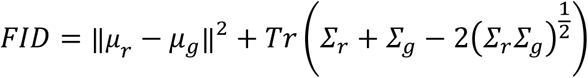

where *μ_r_* and *μ_g_* denote the mean feature vectors of the real and generated image distributions, respectively, while *Σ_r_* and *Σ_g_* represent their corresponding covariance matrices.^18^

### CLIP-Based Binary Classification Network

We utilized the Contrastive Language-Image Pre-training (CLIP) architecture, whose image encoder is based on a Vision Transformer. CLIP is pre-trained on over 400 million image-text pairs, enabling it to learn highly generalized and transferable visual representations. To implement this within our pipeline, we employed OpenCLIP, an open-source PyTorch implementation of the original CLIP model.^11^

We used the ViT-L/14 image encoder with the official OpenAI pre-trained weights. The original 256×256 RGB SEM images were processed using the standard OpenCLIP preprocessing pipeline for the OpenAI-pretrained ViT-L/14 model, which resizes and crops them to the model’s 224×224 input resolution. Instead of training the entire model from scratch, we fine-tuned selected layers of the pre-trained model on the SEM datasets.^11^ For each dataset, the authentic and synthetic SEM images were divided into training and test sets using a 70:30 split. The training set was used to fine-tune the CLIP-based classifier, while the test set was kept separate and used only for evaluating its performance.

Most of the network parameters were frozen to retain the pre-trained representations. The final two residual blocks (*resblocks[-1]* and *resblocks[-2]*) of the visual transformer were unfrozen and fine-tuned on the SEM datasets, allowing these layers to adapt to the characteristics of the target images while the remaining parameters were kept fixed.^11,22^

The CLIP image embedding is passed through a custom binary classification head. During the forward pass, the encoded image features extracted by the ViT-L/14 backbone undergo L2 normalization along the embedding dimension.^11,22^ These normalized features are then passed through a Multi-Layer Perceptron (MLP) structured as follows:

- A linear transformation layer that projects the CLIP embedding dimension down to a 256-dimensional latent space.
- A Rectified Linear Unit (ReLU) activation function to introduce non-linearity.
- A Dropout layer with a probability of 0.3 to mitigate overfitting during the training process.
- A final linear projection that outputs a single scalar logit for binary classification.

For network optimization, we utilized the AdamW optimizer with a learning rate of 1 × 10^−5^ and a weight decay coefficient of 1 × 10^−4^. Parameter updates were applied exclusively to the unfrozen transformer blocks and the custom MLP parameters. The loss was calculated using Binary Cross-Entropy with Logits (BCEWithLogitsLoss), which combines the sigmoid activation and binary cross-entropy loss into a single numerically stable operation. The network was trained for 15 epochs using a batch size of 32.^11,22^

Classifier performance was evaluated using accuracy and Average Precision (AP). Accuracy measures the proportion of images correctly classified as authentic or synthetic using a probability threshold of 0.5 and was calculated as

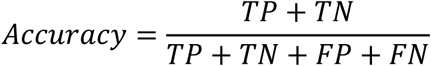

where *TP*, *TN*, *FP*, and *FN* denote the numbers of true positives, true negatives, false positives, and false negatives, respectively.^23^ AP was calculated from the continuous prediction scores produced by the classifier and summarizes the precision–recall relationship across different classification thresholds. It was calculated as

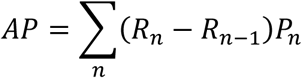

where *P_n_* and *R_n_* are the precision and recall at the *n*-th prediction threshold, respectively.^24^ Both metrics were reported to provide complementary measures of classification performance: accuracy evaluates the final binary predictions at the selected threshold, whereas AP evaluates the quality of the continuous prediction scores across thresholds.^23,24^

### Visual Interpretability Using EigenCAM

We integrated EigenCAM (Eigen-Class Activation Mapping) to generate localized, visual explanations of the model’s decision-making process. EigenCAM calculates the first principal component of the 2D activations from the target convolutional or transformer layer.^19^

Our implementation utilized the *PyTorch Grad-CAM* library.^25^ To capture the most mature, high-level feature representations before classification, we designated the final residual block of the CLIP ViT-L-14 backbone (*model.clip.visual.transformer.resblocks[-1]*) as the target layer, since it provides the final high-level visual representations before the image embedding is passed to the classification head.^19,25^

Vision Transformers represent images as a sequence of image patches.^11^ To visualize these patch-level activations as a 2D heatmap, we used a custom *reshape_transform* function. The function removed the initial Class (CLS) token from the 257-token output of the final transformer block and reshaped the remaining 256 image tokens into a 16×16 spatial grid. The resulting dimensions are then rearranged to standard image format *[Batch, Channels, Height, Width]*, restoring the spatial context of the extracted features.^22^ Since the *PyTorch Grad-CAM* library expects standard multi-class outputs,^25^ we encapsulated the binary CLIP classifier within a custom *CAMWrapper*. This wrapper intercepts the single 1D logit output and introduces an auxiliary dimension (*unsqueeze(1)*), ensuring compatibility with the EigenCAM pipeline.^19,25^

During inference, the preprocessed image was passed through the classifier to obtain a logit, which was converted to a probability using the sigmoid function. A probability greater than 0.5 was assigned to the “Fake” class. In parallel, EigenCAM was used to generate an activation map from the selected transformer layer. The resulting map was upsampled and overlaid on the corresponding RGB image, resized to 224×224 pixels, to visualize the image regions associated with strong activations in the selected layer.^19,25^

## RESULTS AND DISCUSSION

The nanoparticle SEM dataset was obtained from the publicly available dataset reported by Boiko *et al.*,^26^ which contains SEM images of a wide variety of nanoparticle morphologies (**Figure 1A**). The biological SEM dataset was compiled from publicly available SEM images of biological specimens reported by Aversa *et al.*^27^ (**Figure 1B**). The nanoparticle and the biological datasets consisted of 750 and 953 authentic SEM images, respectively, representing the “real” class.^26,27^

**Figure 1:**
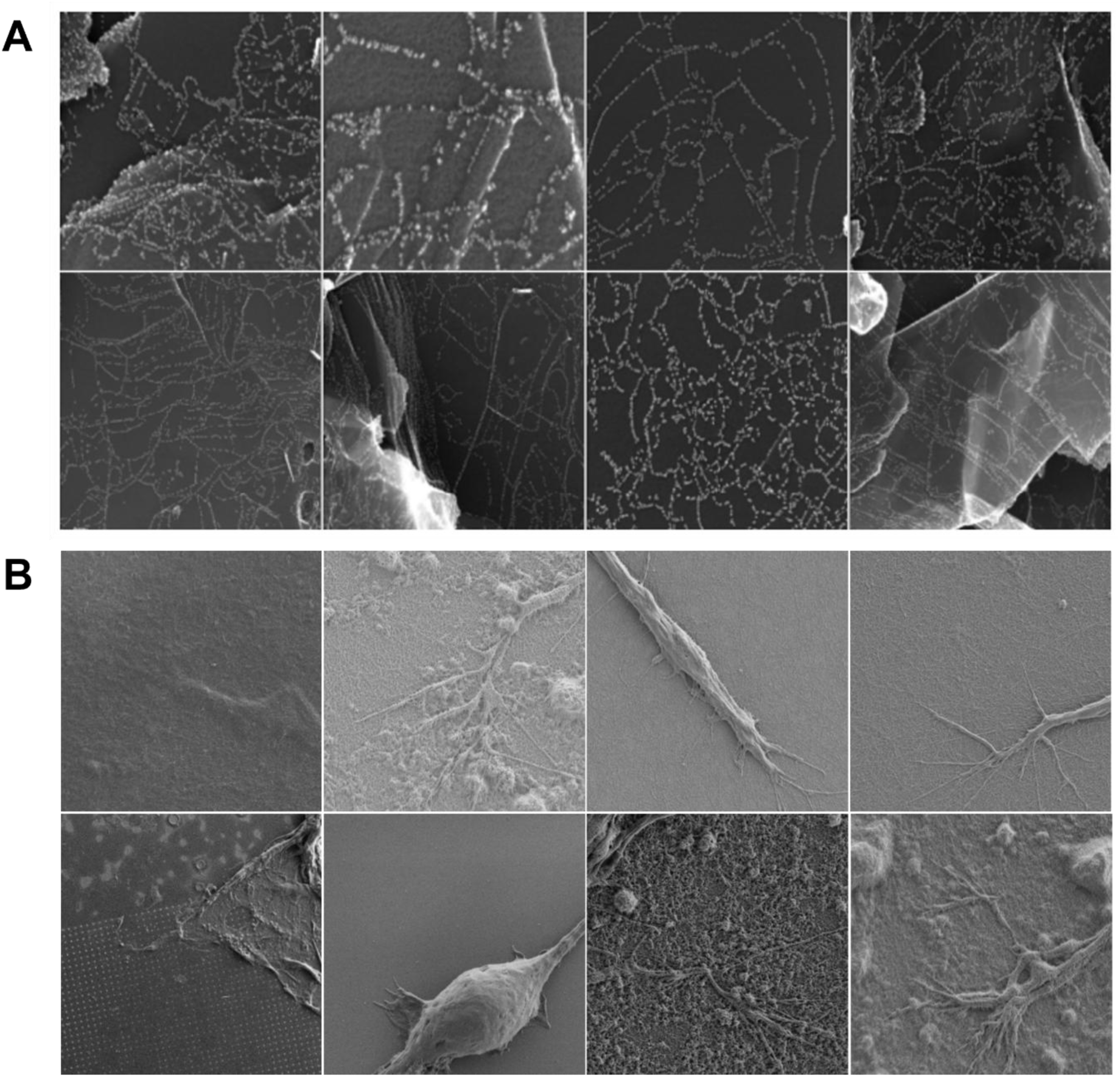
Real SEM sample images. The images illustrate the diversity of surface morphologies and contrast present in the real **(A)** nanoparticle and **(B)** biological SEM datasets used for training and evaluation.

FastGAN was first evaluated for its ability to generate synthetic SEM images that resemble those in the real datasets. Representative images generated for the nanoparticle and biological datasets are shown in **Figure 2A** and **Figure 2B**, respectively. We computed the Mean Squared Error (MSE) distance between each real image and the set of generated images, plotting the minimum distance observed for each real image. This process enabled us to evaluate the pixel-wise similarity between the generated and real images, as illustrated in **Figure 2C** and **Figure 2D** for the nanoparticle and biological datasets, respectively.

**Figure 2:**
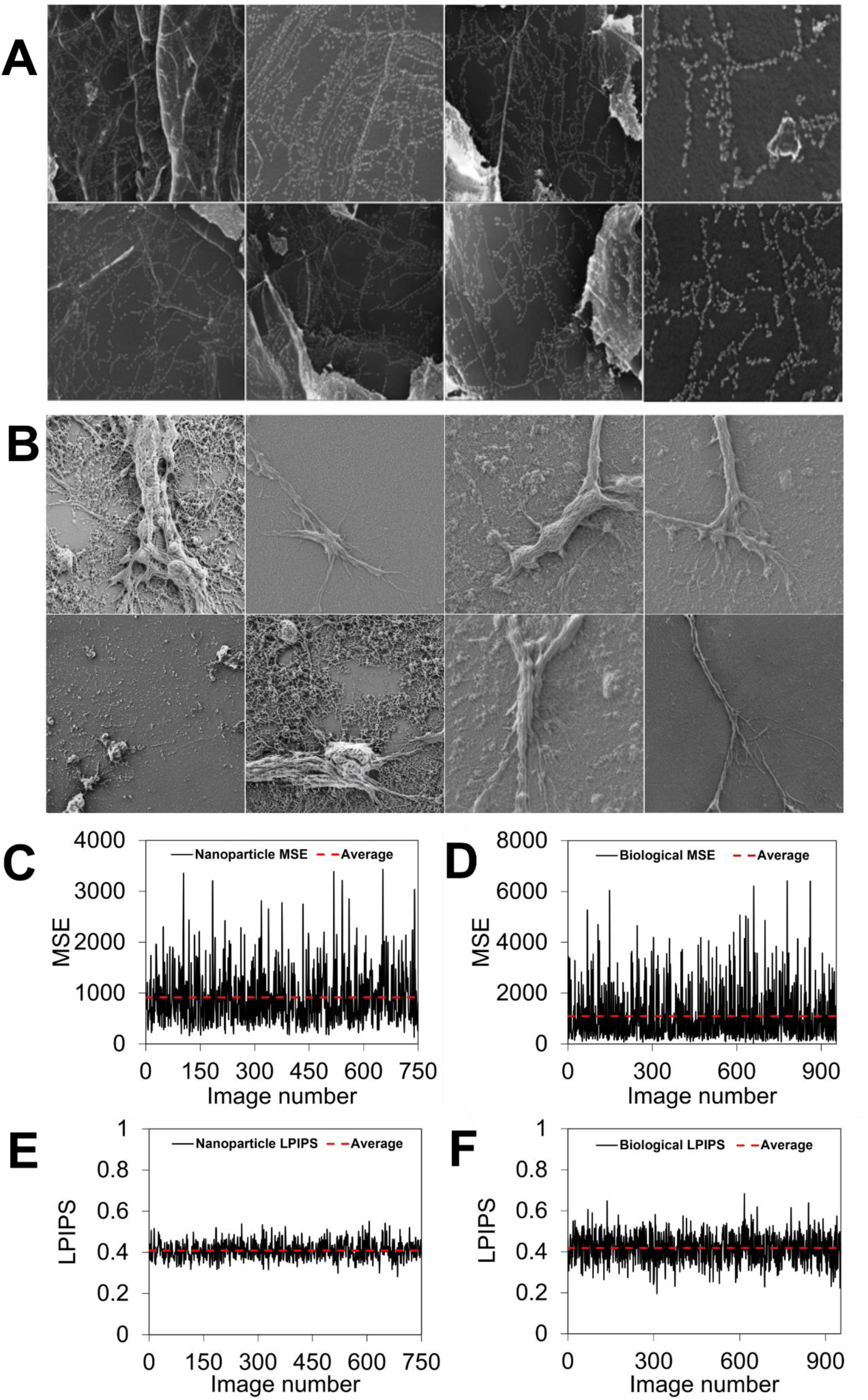
Quantitative evaluation of FastGAN-generated SEM images. **(A)** Sample-generated nanoparticle SEM images. **(B)** Sample-generated biological SEM images. **(C)** Mean Squared Error (MSE) distances for the nanoparticle dataset. **(D)** MSE distances for the biological dataset. **(E)** Learned Perceptual Image Patch Similarity (LPIPS) distances for the nanoparticle dataset. **(F)** LPIPS distances for the biological dataset. For both datasets, the average MSE and LPIPS values are shown as red dashed lines.

For the nanoparticle dataset, the minimum MSE distance recorded was around 183, while the average MSE was 913.05±548.78. For the biological dataset, the minimum MSE distance was around 87, with an average MSE of 1088.86±1025.32. The lower minimum MSE for the biological dataset indicates that at least some generated images were more closely matched to individual real images at the pixel level. However, its higher average MSE indicates that the generated image closest to the corresponding real image differed more from it than in the nanoparticle dataset.^17^ In both datasets, the nonzero minimum MSE values indicate that none of the generated images was pixel-identical to the real images used for comparison.

For each real image, we computed the LPIPS distance to all generated images and plotted the minimum distance observed in **Figure 2E** for the nanoparticle dataset and **Figure 2F** for the biological dataset. For the nanoparticle dataset, the minimum LPIPS value was 0.28, with an average LPIPS of 0.41±0.04. For the biological dataset, the minimum and average LPIPS values were 0.20 and 0.42±0.07, respectively. The similar mean LPIPS values indicate that, on average, the generated images closest to the real SEM images exhibited comparable perceptual resemblance across both datasets, based on their learned visual features. The lower minimum value for the biological dataset indicates that at least one generated image had a smaller perceptual distance from a real image. Additionally, the non-zero minimum LPIPS values indicate that none of the generated images was perceptually identical to the real images under the LPIPS metric.^14^

To further evaluate the similarity between the generated and real images at the distribution level, the Fréchet Inception Distance (FID) was computed between the real and FastGAN-generated image sets for both datasets. The nanoparticle dataset yielded an FID score of approximately 36, whereas the biological dataset yielded an FID score of around 65. Since lower FID values indicate greater similarity between the feature distributions of real and generated images,^18^ the lower FID for the nanoparticle dataset indicates greater distributional similarity between the generated and real nanoparticle images. In contrast, the higher FID for the biological dataset indicates greater distributional differences between the generated and real biological images. Together with the MSE and LPIPS results, these findings show that FastGAN-generated images exhibit measurable similarity to the corresponding real SEM image distributions, with the degree of similarity varying between the two datasets.

We next evaluated images generated using StyleGAN2-ADA for the nanoparticle and biological SEM datasets. Representative generated images are shown in **Figure 3A** for the nanoparticle dataset and **Figure 3B** for the biological dataset. The quantitative evaluation, shown in **Figures 3C–3F**, demonstrates trends similar to those observed for FastGAN. For the nanoparticle dataset, the minimum and average MSE values were 199 and 956.99±544.24, respectively, while the biological dataset yielded a minimum MSE of 97 and an average of 1079.65±1027.19. As observed with FastGAN, the biological dataset exhibited a lower minimum MSE but a higher average MSE, indicating that some generated biological images were more closely matched to individual real images at the pixel level, while the closest generated image was, on average, more different from the real images than in the nanoparticle dataset. In both datasets, the non-zero minimum MSE values indicate that none of the generated images was pixel-identical to the real images used for comparison.

**Figure 3:**
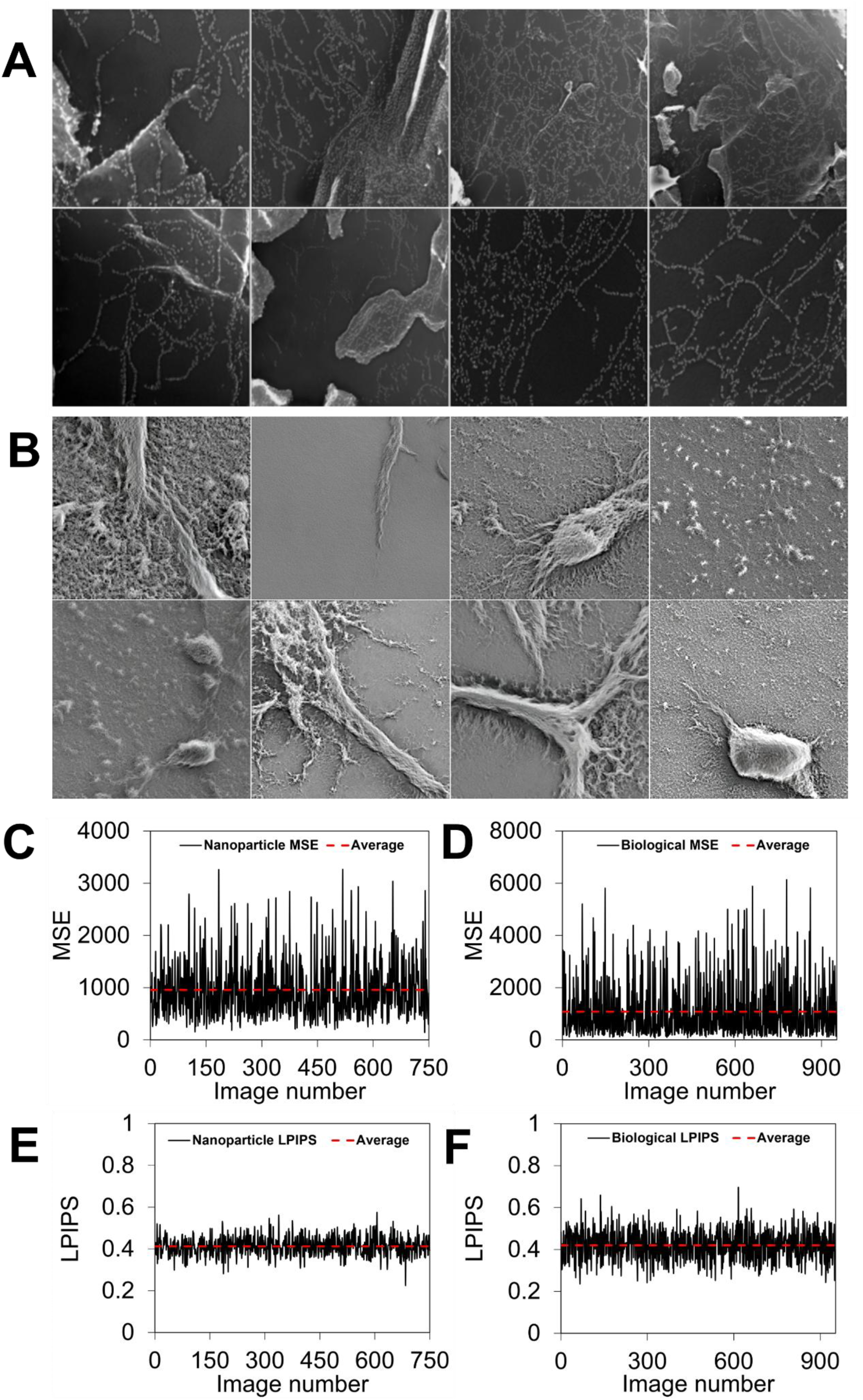
Quantitative analysis of images generated using StyleGAN2-ADA. **(A)** Sample-generated nanoparticle SEM images. **(B)** Sample-generated biological SEM images. **(C)** Mean Squared Error (MSE) distances for the nanoparticle dataset. **(D)** MSE distances for the biological dataset. **(E)** Learned Perceptual Image Patch Similarity (LPIPS) distances for the nanoparticle dataset. **(F)** LPIPS distances for the biological dataset. For both datasets, the average MSE and LPIPS values are shown as red dashed lines.

The LPIPS analysis showed similar perceptual distances between the generated and real images across the two datasets. For the nanoparticle dataset, the minimum and average LPIPS values were 0.22 and 0.41±0.04, respectively, whereas the biological dataset achieved 0.24 and 0.42±0.07, respectively. The similar mean minimum LPIPS values indicate that the generated images with the lowest perceptual distances exhibited comparable perceptual resemblance to the real images across the two datasets.

The nanoparticle and biological datasets achieved FID scores of approximately 44 and 106, respectively, and the higher FID for the biological dataset indicates a greater difference between the generated and real biological image distributions.^18^ Compared with FastGAN, StyleGAN2-ADA produced higher FID values for both datasets, indicating that FastGAN more closely matched the feature distributions of the corresponding real SEM images under the conditions evaluated. Taken together with the MSE and LPIPS results, the FID results indicate that StyleGAN2-ADA-generated images exhibited measurable similarity to the corresponding real SEM image distributions, although the degree of similarity varied across datasets and evaluation metrics. The relatively large standard deviations for the biological dataset indicate that both the generative models did not reproduce all real images to the same degree of pixel-level similarity, suggesting that some of the generated images matched some biological morphologies more closely than others.

The ability of the fine-tuned CLIP classifier to distinguish authentic SEM images from GAN-generated images was evaluated across the nanoparticle and biological datasets, and the classification results are summarized in **Table 1**. For the nanoparticle classifier, the overall test accuracy and AP were 97% and 99.2%, respectively. When evaluated separately on the generator-specific test sets within the dataset, the classifier achieved 95.3% accuracy and 98.5% AP on FastGAN, and 98.7% accuracy and 99.9% AP on StyleGAN2-ADA. The biological classifier achieved overall accuracy and AP of 99.8% and 100%, respectively, with 100% accuracy and AP on FastGAN, and 99.6% accuracy and 100% AP on StyleGAN2-ADA.

**Table 1:**
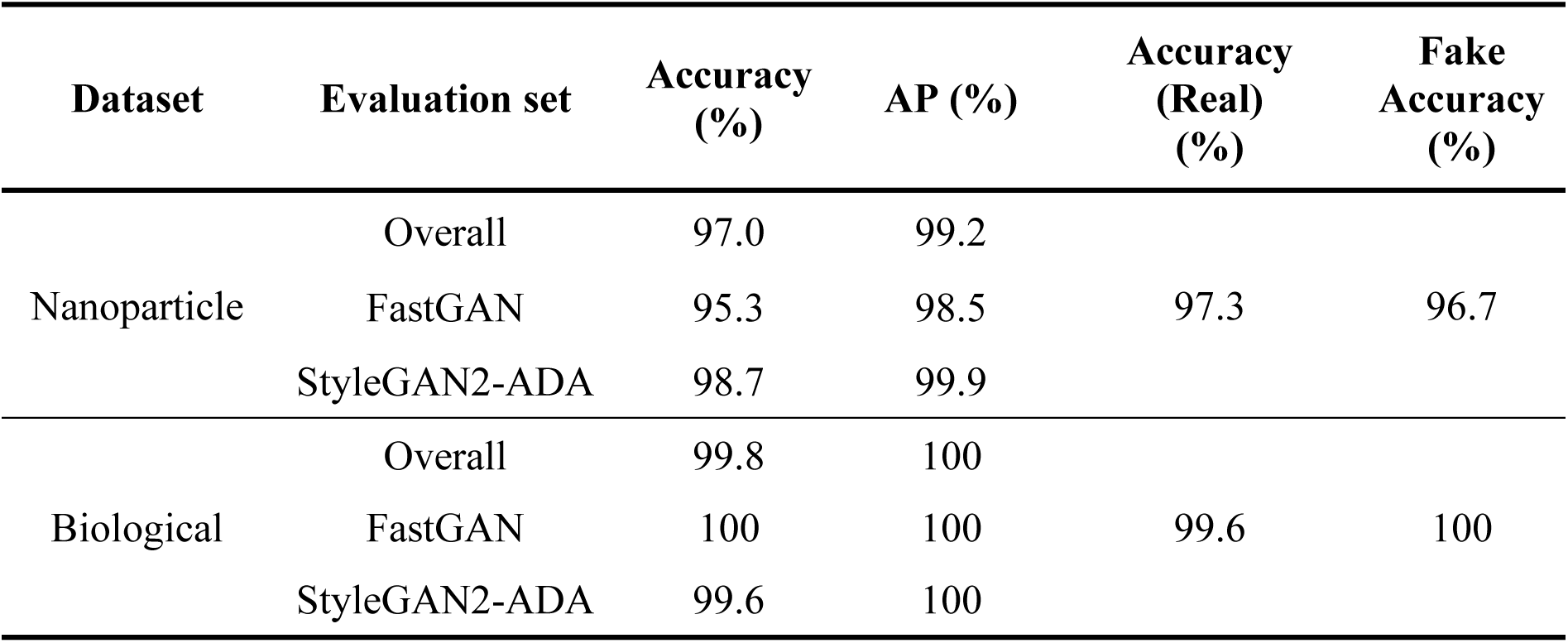
Classification performance of the proposed CLIP-based classifier on nanoparticle and biological SEM datasets generated using StyleGAN2-ADA and FastGAN.

The classifier maintained high classification performance across both SEM datasets and generative architectures. The differences in accuracy between StyleGAN2-ADA and FastGAN varied across the two datasets, with FastGAN showing lower accuracy on the nanoparticle dataset but a higher accuracy on the biological dataset. The corresponding AP values remained high across all evaluated conditions, indicating that the classifier produced well-separated prediction scores for authentic and synthetic images across the tested GAN architectures. However, these results represent evaluations within the respective training domains and therefore do not by themselves establish cross-domain generalization.

To assess the transferability of the proposed framework across SEM imaging domains, additional cross-domain experiments were performed. For that, the classifier trained on the nanoparticle dataset was applied to the biological test set, while the classifier trained on the biological dataset was applied to the nanoparticle test set. For each evaluation, the complete target-domain test set containing authentic images and images generated by both GAN architectures was used. Performance decreased during cross-domain evaluation compared with within-domain testing, as depicted in **Table 2**, reflecting the substantial morphological differences between the two datasets. The nanoparticle-trained classifier achieved 66.6% accuracy and 88.6% AP on the biological dataset, whereas the biological-trained classifier achieved 73.6% accuracy and 87.0% AP on the nanoparticle dataset. The reduction in accuracy compared with the corresponding within-domain evaluations indicates that changes in SEM morphology and image characteristics affect the classifier’s performance when applied to a different domain. However, the relatively high AP values indicate that the prediction scores retained useful separation between authentic and synthetic images under this domain shift. These findings indicate that while domain-specific training provides the highest detection accuracy, the proposed CLIP-based framework exhibits encouraging cross-domain generalization capability. Further improvements in cross-domain performance may be achieved by training on more diverse SEM datasets encompassing a broader range of material classes and morphologies.

**Table 2:** Cross-domain performance of the dataset-specific CLIP classifiers on nanoparticle and biological SEM datasets.

| Training Domain | Testing Domain | Accuracy (%) | AP (%) |
| --- | --- | --- | --- |
| Nanoparticle | Biological | 66.6 | 88.6 |
| Biological | Nanoparticle | 73.6 | 87.0 |

Although the fine-tuned CLIP model achieves a certain level of classification performance, the basis of its predictions is not directly interpretable. In materials science, examining the image regions that contribute to the model’s predictions is important for assessing whether the classifier relies on meaningful image features rather than spurious visual patterns or background information.^28^

To understand how the fine-tuned CLIP model makes its decisions, we used EigenCAM to qualitatively examine the image regions that contribute to the feature representations learned by the proposed classifier.^19,25^ Representative EigenCAM overlays from both the nanoparticle and biological datasets are shown in **Figure 4**. By overlaying these heatmaps onto the input images, we can visualize the regions of the images that contribute most strongly to the learned feature representations. The visualizations were generated from the final transformer block of the CLIP ViT-L/14 image encoder, as this layer contains the high-level visual representations immediately before image embeddings are used for classification.^11,22^

**Figure 4.**
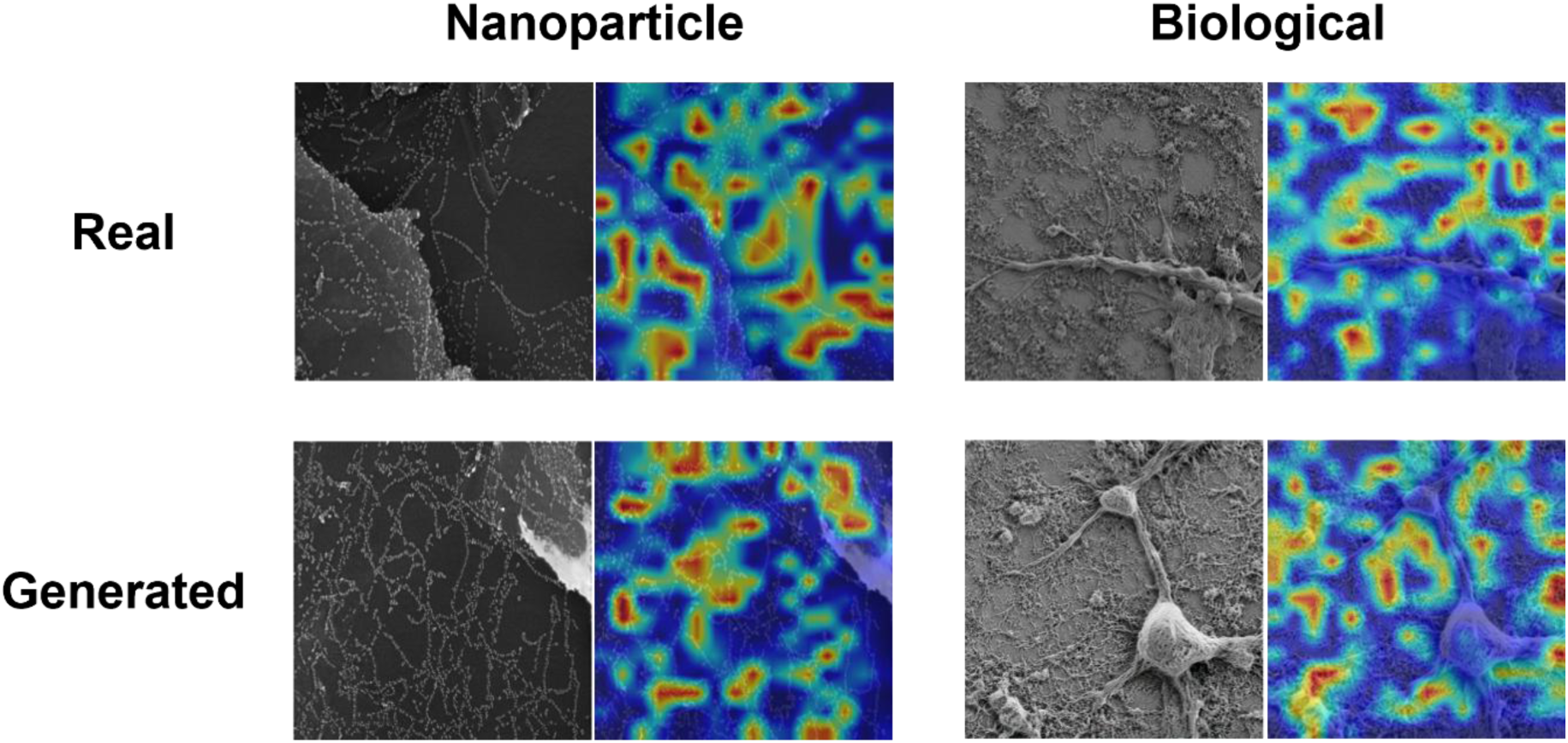
Representative SEM images and their corresponding EigenCAM overlays for real and GAN-generated images from the nanoparticle and biological datasets. The EigenCAM visualizations were generated from the final transformer block of the fine-tuned CLIP ViT-L/14 image encoder and highlight the image regions contributing most strongly to the learned feature representations used for classification.

A noticeable observation is that many activation regions appear as square or block-like patches. This behavior is expected because the CLIP image encoder is based on a Vision Transformer (ViT), which processes an image by dividing it into fixed-size patches. Consequently, the EigenCAM visualization reflects the model’s patch-based feature extraction, resulting in activations that often follow the boundaries of these image patches.^11^

No obvious qualitative differences were observed between the EigenCAM overlays of real and fake SEM images. The activation patterns appeared broadly similar across the two classes, with strong activations distributed across multiple regions of the SEM images rather than consistently concentrated in a single localized area.

It is important to note that EigenCAM visualizes regions associated with strong activation patterns in the selected feature representations and does not directly provide class-specific attribution for the real/fake prediction.^19^ Therefore, the similar activation patterns observed for both classes do not contradict the high classification accuracy achieved by the proposed model. Rather, they indicate that similar image regions are strongly represented in the selected transformer layer for both real and synthetic SEM images.

Although the proposed classifier demonstrated strong performance on the SEM datasets considered in this study, the images were collected from a limited number of material categories. SEM imagery encompasses a wide range of application domains, including nanoparticles, biological cells, nanoplates, nanowires, porous materials, and numerous other micro- and nanostructures, each exhibiting distinct morphological characteristics.^29^ Training a single generative model on a highly heterogeneous collection of such images may increase the complexity of the underlying data distribution, potentially making image generation more challenging, particularly when only limited training data are available.^15,30^ Consequently, a direction for future work would be to investigate category-specific generative models and corresponding specialized detection models for individual SEM domains. Such an approach may improve both the fidelity of generated images and the robustness of fake-image detection by allowing the models to learn more homogeneous structural and textural characteristics. Furthermore, evaluating the proposed CLIP-based classifier across a broader range of SEM categories and previously unseen material systems would provide a more comprehensive assessment of its generalization capability.

## CONCLUSION

In this work, we present a deep learning framework utilizing a Contrastive Language-Image Pre-training (CLIP) Vision Transformer (ViT-L/14) backbone to identify synthetic Scanning Electron Microscopy (SEM) images generated using FastGAN and StyleGAN2-ADA. As generative architectures become increasingly capable of producing hyper-realistic micrographs, safeguarding scientific integrity in materials science necessitates highly reliable, automated verification tools^10^. The classifier achieved accuracies ranging from 95% to 100% across the nanoparticle and biological datasets when evaluated within their respective domains. We also generated synthetic SEM images using FastGAN and StyleGAN2-ADA and evaluated their similarity to real SEM images using MSE, LPIPS, and FID, revealing differences in image similarity across datasets and generative architectures. Cross-domain evaluation showed that the classifier retained discriminative information when applied to a different SEM dataset, although classification accuracy decreased under domain shift. EigenCAM visualizations further showed that strong activations were distributed across different regions of the SEM images rather than being concentrated in a single localized area. Overall, this study demonstrates that a CLIP-based vision foundation model can detect GAN-generated SEM images across different generative architectures and SEM domains, while also highlighting the effects of domain differences on detection performance.

## ACKNOWLEDGEMENTS

The authors would like to sincerely thank IIT Gandhinagar for providing the facilities and resources necessary to carry out the study, especially the library facility for research paper curation and PARAM Ananta (supercomputer) for carrying out the model training.

## CODE AND DATA AVAILABILITY

All the pre-processed datasets, the classifier models and their corresponding codes are freely available at https://github.com/adwaith19/SEM-Detection.

## AUTHOR CONTRIBUTIONS

A. Palangattu led the study, performed the computational work and analysis, and drafted the manuscript. A. K. Sah curated and pre-processed the datasets. K. Pushpavanam provided primary supervision, guided the research design and interpretation of the results, and S. Raman critically reviewed the manuscript. All authors reviewed and approved the final manuscript.

